# Quantitative modeling of fine-scale variations in the *Arabidopsis thaliana* crossover landscape

**DOI:** 10.1101/2021.10.06.463263

**Authors:** Yu-Ming Hsu, Matthieu Falque, Olivier C. Martin

**Affiliations:** Université Paris-Saclay, CNRS, INRAE, Univ Evry, Institute of Plant Sciences Paris-Saclay (IPS2), 91405 Orsay, France; Université de Paris, CNRS, INRAE, Institute of Plant Sciences Paris-Saclay (IPS2), 91405 Orsay, France; Université Paris-Saclay, INRAE, CNRS, AgroParisTech, GQE - Le Moulon, 91190 Gif-sur-Yvette, France

## Abstract

In essentially all species where meiotic crossovers have been studied, they occur preferentially in open chromatin, typically near gene promoters and to a lesser extent at the end of genes. Here, in the case of *Arabidopsis thaliana*, we unveil further trends arising when one considers contextual information, namely summarized epigenetic status, size of underlying genomic regions and degree of divergence between homologs. For instance we find that intergenic recombination rate is reduced if those regions are less than 1.5 kb in size. Furthermore, we propose that the presence of single nucleotide polymorphisms is a factor driving enhanced crossover rate compared to when homologous sequences are identical, in agreement with previous works comparing rates in homozygous and heterozygous blocks. Lastly, by integrating these different factors, we produce a quantitative and predictive model of the recombination landscape that reproduces much of the experimental variation.

## Introduction

Crossovers formed during meiosis drive the shuffling of allelic combinations when going from one generation to the next. They thereby play a central role in genetics and evolution and they are also key in all forms of breeding. In cultivated plants, the pericentromeric regions tend to be large and refractory to crossovers (Bauer *et al*., 2013; Choulet *et al*., 2014). Although these regions have a high density of transposable elements, in crops they nevertheless contain a sizable number of genes. Attracting crossovers (COs) into these regions could have benefits for genetic studies (e.g., to identify gene functions) and for selection of new combinations of alleles of relevance for breeding.

CO formation processes (Villeneuve *et al*., 2001; Mercier *et al*., 2015) start with the active formation of double strand breaks (Keeney & Neale, 2006) and end with DNA repair, leading to either CO or non-COs (Hunter, 2015). They are tightly regulated, in particular they ensure at least one CO per bivalent (Jones & Franklin, 2006; Zickler & Kleckner, 2016) but not many more in spite of huge variations in genome size (Fernandes *et al*., 2018). Furthermore, CO *distribution* tends to be very heterogeneous *along* chromosomes, indicating that there are also determinants of CO formation at finer scales. Typically, pericentromeres and more generally regions rich in heterochromatin are depleted in COs. In contrast, regions of open chromatin such as gene promoters are enriched in COs. In yeast it has been possible to measure precisely the distribution of double strand breaks (precursors of both COs and non-COs), revealing a very high level of heterogeneity genome-wide (Pan *et al*., 2011). It is generally assumed that such heterogeneities, detected all the way down to the scale of a few kb, arise also for CO distributions in most species, though the resolution of CO maps have not been able to fully confirm this expectation. That is particularly true in plants that are our focus here and for which the best dataset averages about one CO every 3.5 kb (Rowan *et al*., 2019).

Our objective here is to shed light on genomic and epigenomic features that shape recombination rate on fine scales in *A. thaliana*, a species chosen because it has more extensive CO datasets than other species. Nevertheless, the data are not totally adequate. Firstly, high resolution CO maps are sex-averaged because produced using F_2_ populations, so we cannot address sex-dependencies of crossover landscapes. Secondly, although it is widely accepted that open chromatin favors CO formation, we lack information on the state of chromatin during meiosis, *i*.*e*., in the actual cells (meiocytes) where COs arise. In the absence of such data, we have to rely on epigenetic measurements in other types of cells, germinal or somatic. Modulo these caveats, we exploit a recent high resolution dataset detecting 17077 COs in a large *A. thaliana* F_2_ population (Rowan *et al*., 2019). Analysis of these COs provides new insights. For instance, the *size* of an intergenic region affects recombination rate, there being a suppression for regions whose size is less than about 1.5 kb. Furthermore, it is likely that COs are partly suppressed by lack of SNPs, a result that explains the «heterozygous block effect» found previously (Ziolkowski *et al*., 2015) whereby the insertion of a heterozygous block into an otherwise homozygous region enhances recombination rate therein. Use of additional datasets (Blackwell *et al*., 2020) provides further evidence for this suppression effect. These different insights allow us to build a quantitative model that integrates genomic information, local epigenetic status and contextual factors. This model has good predictive power and reproduces much of the recombination rate variation in *A. thaliana*, pointing to the importance of different contextual factors modulating local crossover rate.

## Materials and Methods

### CO datasets

COs are inferred to lie within intervals, delimited by SNPs, anchoring transitions between homozygous and heterozygous regions (Rowan *et al*., 2019). When measuring recombination rate in a given bin, we count one CO for each CO interval lying completely within that region, and otherwise we apply the simple *pro-rata* rule. We downloaded the dataset of CO intervals of Rowan *et al*. (Rowan *et al*., 2019) based on 2182 F_2_ individuals from a cross between Col-0 and Ler. We also used the data of 5 F_2_ populations based on crossing Col-0 with other 5 accessions (Blackwell *et al*. 2020). The associated files were kindly provided by Ian Henderson, University of Cambridge, Cambridge, UK, and are included as Supplementary Material. For the whole study, the experimental recombination rate *r* (in cM/Mb) was calculated using the formula: *r =* 100 * *n*_*co*_ */(n*_*plant*_ *\* 2 * L*_*Mb*_) where *n*_*co*_ is the number of crossover intervals contained in the relevant bin or region, *n*_*plant*_ is the number of F_2_ plants, and is the length of the bin in Mb.

### Genomic annotation of Col-0 and structural variations between Col-0 and Ler genomes

For Col-0 genomic features, we utilized TAIR10 annotation specifying coding genes and super families of transposable elements. We compared the TAIR10 reference Col-0 genome and the Ler assembled genome to detect syntenic regions and structural variations (SVs) (Berardini *et al*., 2015; Jiao & Schneeberger, 2020). SVs were identified using MuMmer4 and SyRI software (Goel *et al*., 2019). The parameters used in the “nucmer” function of MuMmer4 were set via “-l 40 -g 90 -b 100 -c 200”. All genomic and epigenomic features were computed after masking out the regions containing the structural variations defined by SyRI.

### Col-0 epigenomic features and segmentation of chromosomes into chromatin states

BigWig, bedGraph and bed files of H3K4me1, H3K4me3, H3K9me2, H3K27me3, ATAC and DNase measurements on Col-0 were downloaded from the NCBI and ArrayExpress databases (cf. Supplementary Table S1). Segmentation of the chromosomes into 9 chromatin states were obtained from the study of Sequeira-Mendes *et al*. (2014) which again is specific to Col-0.

### Identifying SNPs in the 5 F2 populations

The 5 F_2_ populations (Blackwell *et al*., 2020) had Col-0 as shared parent, the other parent was Ler, Ws, Ct, Bur or Clc. Their sequences were downloaded from the ArrayExpress database (accession identifiers E-MTAB-5476, E-MTAB-6577, E-MTAB-8099, E-MTAB-8252, E-MTAB-8715, E-MTAB-9369). For aligning the reads to the TAIR10 reference genome (Berardini *et al*., 2015), we used the “mem” algorithm of Burrows-Wheeler Alignment (BWA-MEM; v0.7.17) (Li, 2013), then samtools (v1.10) (Li, 2011) and bcftools (v1.12) for SNPs calling. Finally, we applied filters to keep SNPs with (1) a quality score ≥ 100, (2) mapping quality score ≥ 20, (3) depth below 2.5 mean depth of the corresponding F_2_ population to eliminate anomalously high coverages indicative of multi mappings, and (4) positions that only contained uniquely mapped reads.

### Testing causality of the SNP density effect by comparing H0 and H1

Under H0 and H1 there is an underlying “reference” recombination landscape common to all crosses. H1 allows for a causal SNP density effect by modulating this bin-dependent reference recombination rate by a multiplicative factor, parametrized as (1 + *α*_1_**ρ**) exp(-*α*_2_**ρ**) where **ρ** is the SNP density for that population and bin while α_1_ and α_2_ are parameters to be adjusted using the whole dataset (containing all bins and all 5 of the F_2_ populations). In contrast, H0 has no such modulation effect. We defined a chi-square

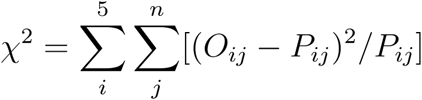

where *O*_*ij*_ and *P*_*ij*_ are the number of observed and H0- or H1-predicted crossovers in the jth bin belonging to the i’th F_2_ population, respectively. This chi-square is minimized to infer α_1_ and α_2_ in addition to the reference recombination rates in each bin. Our test compares the values of chi-square under H1 and H0. To extract an associated *p*-value, we disassociate SNP density and recombination rate by shuffling SNP density between the 5 F_2_ populations, independently in each bin. From this randomization process taking one from H1 to H0, we acquired 1,024 chi-square values that give the distribution of chi square under H0, and then the *p*-value is the probability of having within this distribution a chi square as low as the one of H1. In practice, since high SNP density hinders CO formation (homology is lost at high divergence), we selected the data along the chromosomes where SNP density was in the bottom 50% quantile.

### The quantitative model based on epigenetic states and genomic features

Sequeira-Mendes *et al*. (2014) identified 9 distinct chromatin states in Col-0 segmenting the whole genome. We modified their segmentation as follows. First, noting that heterochromatic regions often contained stretches of alternating states 8 and 9, we relabeled segments of state 8 as state 9 when they were sandwiched between two state-9 segments. This relabeling affected almost exclusively segments in the pericentromeric regions and provided a proxy for heterochromatin. We verified that recombination rate was highly suppressed in such relabeled segments while non relabeled state 8 segments (lying almost exclusively in the arms) did not lead to crossover suppression. Second, we added a new state corresponding to having an SV or insufficient synteny between the two parental genomes of interest.

Given these 10 states and their segmentation of the genome, our model introduces an adjustable “base” recombination rate for each and then applies three modulation factors. The modulation by the intergenic region size is straightforward if one considers a genomic segment lying entirely between two genes; if it does not satisfy that condition, we break it into underlying pieces so that each piece is either entirely within an intergenic region or entirely within a genic region; the modulation is then applied to each piece separately.

The 15 parameters of this quantitative model were identified by fitting to the experimental data using the least square method as the measure of goodness of fit. Specifically, we minimized the sum of squares: 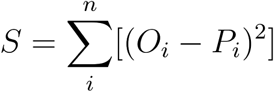 where *O*_*i*_ and *P*_*i*_ represent the number of experimental and predicted crossovers in the i’th bin.

### The software of statistical analysis and visualization

All statistical analysis was based on R 3.63. For fitting model parameters to data, we used the “optim” function with the method “L-BFGS-B”. All visualizations were carried out by the “tidyverse” package (Wickham *et al*., 2019).

## Results

### Standard modeling of crossover rate based on genomic and epigenomic factors is unsatisfactory

Based on 17,077 crossovers from an F_2_ population (Rowan *et al*., 2019), we related recombination rate to the local density of various genomic and epigenomic features. As shown in Fig. 1, the individual relations found are typically non monotonic with correlations of one sign within chromosome arms and of the opposite sign within pericentromeric regions. Such a characteristic makes it difficult to assign a role to any individual feature. This result holds whether using data obtained from somatic tissues or from germinal tissues (cf. Supplementary Fig. S1).

**Figure 1.**
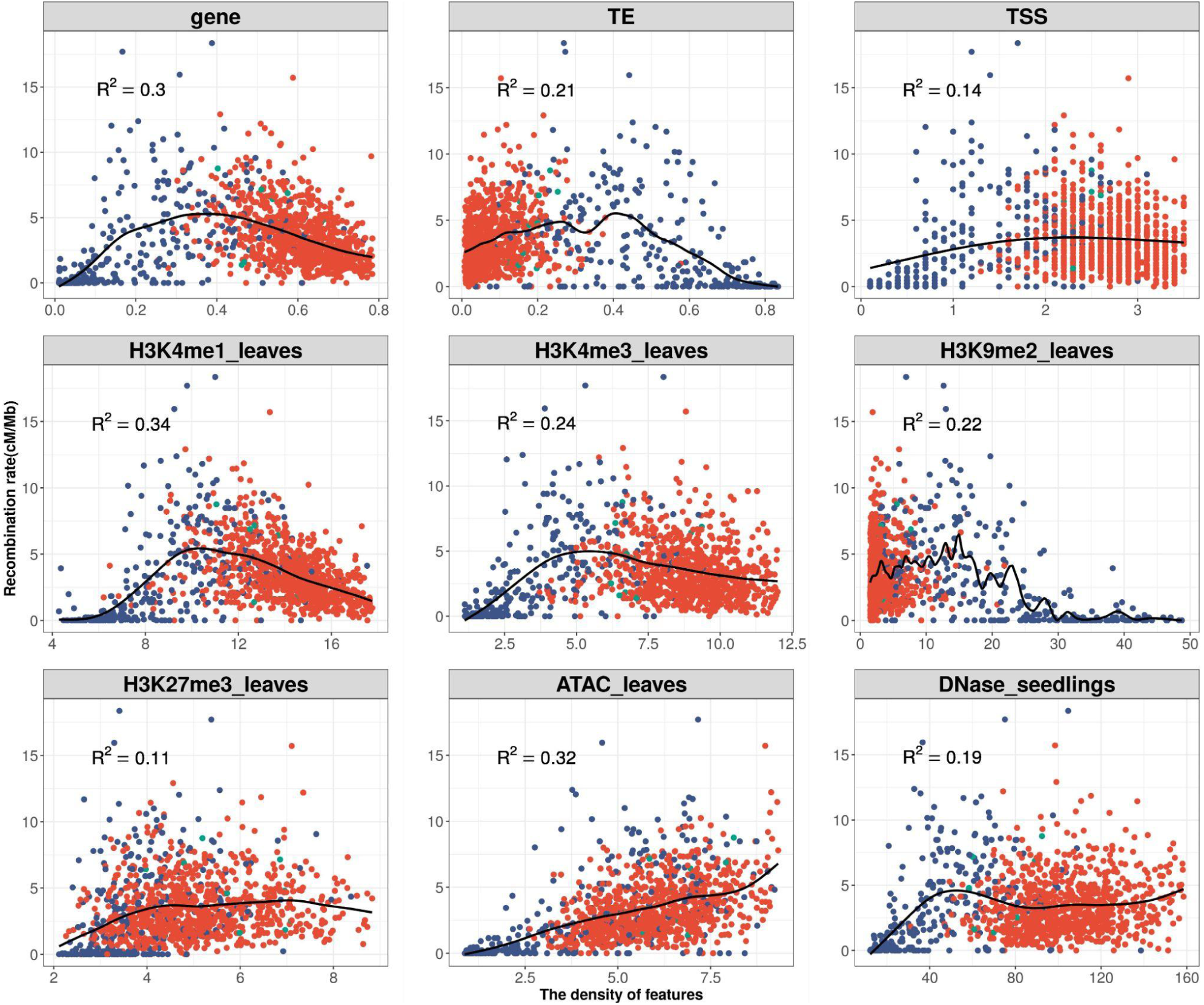
The correlations between recombination rate and 9 genomic or epigenomic features taken from somatic tissues (cf. titles). Each dot represents the values for a 100-kb bin. The x-axis shows the density of each feature, and the y-axis is the recombination rate based on a total of 17,077 crossovers from the Col-0-Ler F_2_ population. Dots in red, blue or green are located in arms, pericentromeric regions or the transition regions between arms and pericentromeric regions, respectively. The fitted black lines were produced by the cubic smoothing spline “smooth.spline” function of the statistical package R. R^2^ corresponds to the fraction of explained variance when using the interpolating splines (Eq. 2). To ensure that the points fill most of the space, the scale is set to display only 95 % of the data, cutting the 2.5 % extremities on both sides of the x-axes in all these plots.

To combine all these factors into a model, the standard approach is to consider an additive framework and then possibly generalize it by including interaction terms. The additive model corresponds to predicting recombination rate within a bin using the following formula:

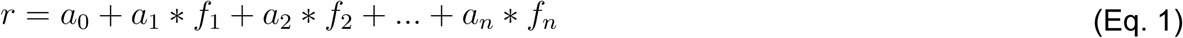

where f_i_ is the density of the ith feature in the bin. In this spirit, we include all 9 feature densities of Fig. 1. In Supplementary Table S2 we provide the fitted values *a*_0_, *a*_1_, …, *a*_9_ when using different bin sizes. Somewhat surprisingly, the coefficient in Eq. 1 for gene coverage density is negative, making the interpretation of the model problematic and suggesting that the additivity assumption is not supported by the data. Finally, to have a measure of goodness of fit, we use the fraction of the recombination rate variation that is “explained” by the model, defined as:

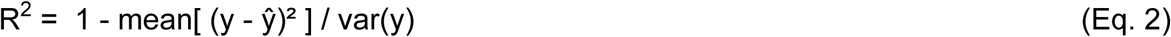

where y is the experimental and ŷ is the predicted value of recombination rate in the different bins along the genome. R^2^ as well as the coefficients in Eq. 1 depend on the bin size; for our “reference” bin size of 100 kb, the model calibration gives R^2^ = 0.36.

To allow for deviations from additivity we follow the standard practice of including interaction terms in the form of pairwise products of feature density values, leading to the formula:

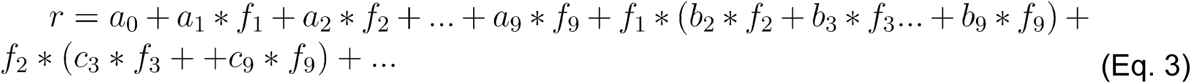

This leads to 46 adjustable parameters *vs* 10 in the additive model. This more complex model explains a fraction R^2^ = 0.35, 0.43, 0.51 and 0.66 of the total recombination rate variance when bin size is 50, 100, 200 and 500kb. Although this is better than the additive model, the interactions do not lead to biological interpretations. Furthermore, the problem of the negative predictions remains, and we also find that the fitted parameters vary substantially with bin size. Thus this model with interactions is not satisfactory and it does not provide insights into the biological determinisms of recombination rate.

### Aggregating genomic and epigenomic features using a chromatin state classifier

Given the drawbacks of the previous modeling framework, we performed aggregation using an automatic classifier approach (Sequeira-Mendes *et al*., 2014), assigning a “chromatin state” to a local region according to a (non linear) combination of such features. The methodology is general but those authors implemented it in the case of Col-0, producing 9 chromatin states based on the combination of 16 genomic or epigenomic features. Their states 8 and 9 correspond to AT-rich and GC-rich heterochromatic regions, respectively, with state 9 being strongly enriched in the pericentromeric regions. Their 7 other states are typically euchromatic. They found that state 1 (respectively state 6) typically colocalizes with transcription start sites (TSS) (respectively transcription termination sites (TTS)). States 3 and 7 are the most abundant states in gene bodies, with the former one tending to be present with state 1 at the 5’ end of genic regions and the latter one arising more frequently in larger transcriptional units. States 2 and 4 typically lie within intergenic regions and they tend to be proximal and distal to the gene’s promoter, respectively. Like state 2 and 4, state 5 is generally within intergenic regions, but it also arises frequently in silenced genes with high levels of H3K27me3.

Because COs form between homologs, we also need to aggregate information about the local synteny between Col-0 and Ler, the two parents of the F_2_ population (Rowan *et al*., 2019) used to estimate the recombination landscape. We thus assign the state “SV” (for Structural Variation) to the non syntenic regions. We then have a total of 10 different “states” that we will study in the rest of this work, referring to them as “chromatin states” even if that is not completely correct. The fraction of the genome covered by any of these chromatin states varies between 5.8% and 13.6%, with state 4 being the most represented and state 8 the least (cf. Fig. 2A, top).

**Figure 2.**
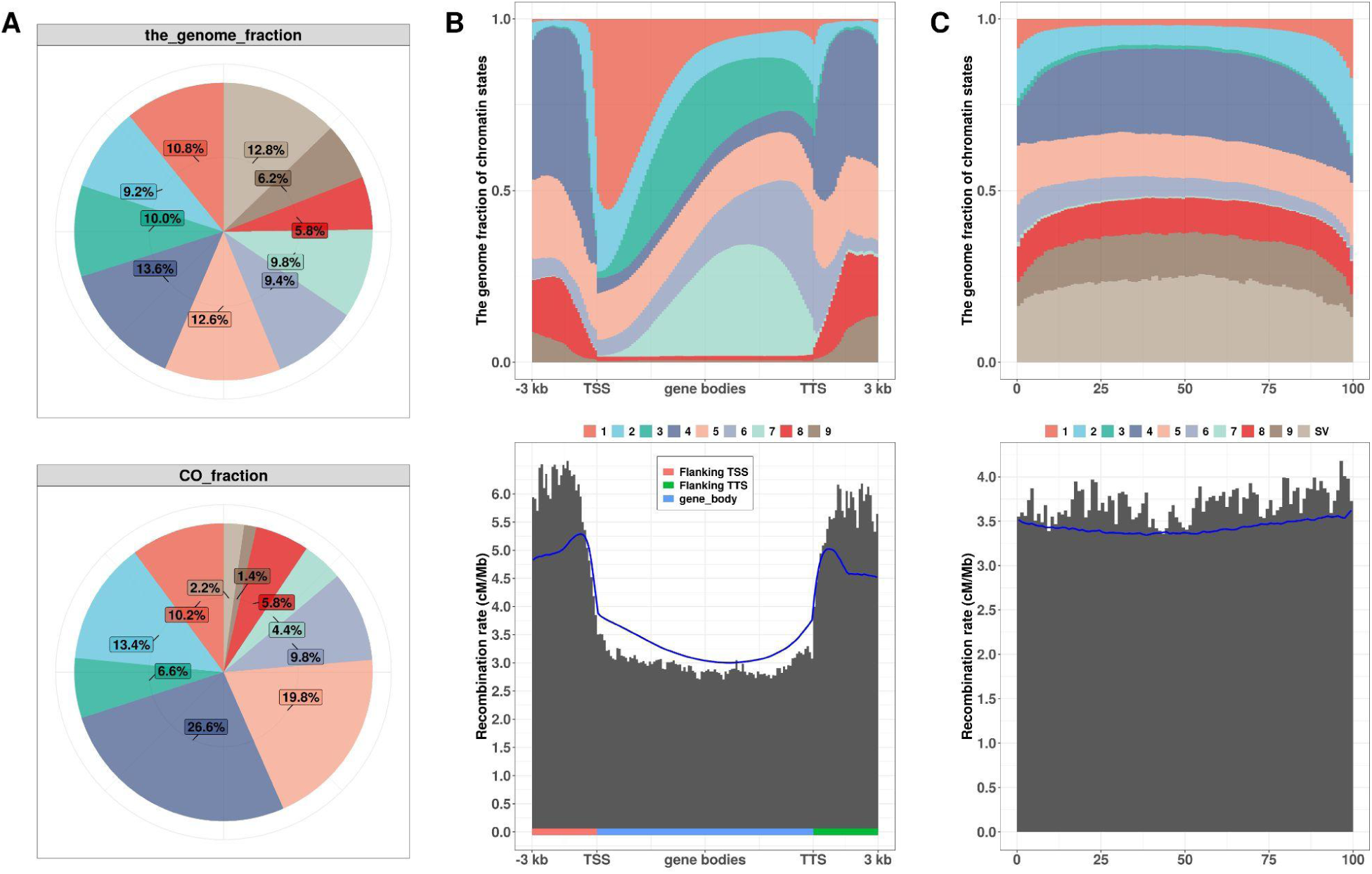
Relations between our 10 chromatin states, genes, intergenic regions, and recombination rate. A) The top pie chart shows the genome-wide occupation percentages of each of the 10 states. “SV” refers to low synteny regions or structural variations between Col-0 and Ler. The lower pie chart shows the percentage of crossover occurrences identified in the 10 states. B) Two plots, giving respectively the profiles of cumulated fractions of occurrences of the 10 different states (top) and the recombination rate pattern (bottom) in cM per Mb, along gene bodies and their 3 kb flanking regions. In the absence of SV, the entire 3-kb flanking region was used, otherwise it was truncated. The gene body goes from the TSS (transcription starting site) to the TTS (transcription termination site) as given in TAIR 10. Only non transposable element coding genes satisfying the synteny filter have been included in the analysis. For the gene body region, the X axis represents *relative* position, that is the distance from the TSS divided by the distance between TTS and TSS. That procedure allows one to pool genes of different sizes. For the flanking regions, X axis represents position relative to the TSS or TTS in kb. The blue curve is the predicted recombination rate when using the chromatin state profiles at the top together with the genome-wide recombination rates derived from A. C) Two plots as in B but now for the intergenic regions. Again the blue curve is the predicted recombination rate when using the chromatin state profiles at the top together with the genome-wide recombination rates derived from A. (The cases of convergent, divergent and parallel transcription have been pooled in this figure, see Supplementary Figs. S4,5,6 for when the three orientations are kept separate.) The legend in the middle of Fig 2B and 2C indicates the corresponding chromatin state of each color used in plotting the chromatin-state profiles.

To transform the trends found by Sequeira-Mendes *et al*. into quantitative patterns we have generated the frequency profiles for each chromatin state as a function of position within gene bodies and their flanking regions. For that task, we used the 25,708 genes extracted from syntenic regions and also considered their extensions on both sides, going out to 3 kb upstream of the TSS and downstream of the TTS. The computed profiles (Fig. 2B (top)) reveal that there is a clear gradient in the chromatin state content along the gene bodies and also along their flanking regions. For instance, the frequency of state 1 has a very sharp rise as one enters the gene on the 5’ side while the frequency of state 7 has a steep fall as one exits the gene on the 3’ side. We performed the analogous computations for intergenic regions and find that the frequency profiles there (cf. Fig. 2C (top)) have much less variation than in gene bodies.

### A simple quantitative model of recombination rate based on discrete chromatin states and structural variations

In contrast to the quantitative factors used in Eq. 1, the state classifier approach identifies discrete states. These can be used as *qualitative factors* in a model of recombination rate by assuming that each state has its own specific recombination rate. This framework both allows for a direct biological interpretation and is mathematically particularly simple. Comparing the genomic fraction of each chromatin state to the observed CO fraction for that state (top and bottom of Fig. 2A) determines the 10 average recombination rates: 3.08, 4.78, 2.16, 6.37, 5.14, 3.48, 1.5, 3.35, 0.7 and 0.57 cM/Mb. Hereafter, these values are referred to as the “experimentally measured state-specific recombination rates”. They are to be compared to the genome-wide average recombination rate of 3.3 cM/Mb. As expected, recombination is strongly suppressed in states 9 and SV.

Second, how well does this “model” predict recombination rates? In Supplementary Fig. S2 we compare experimental and predicted recombination rates when segmenting the genome into bins of size 100 kb. Then the fraction of the variance in the experimental recombination rates that is explained by the model is R^2^ = 0.24. This value is lower than that of the additive model using Eq. 1 (cf. Supplementary Table S2) but note that when using the experimentally measured state-specific recombination rates there are *no adjustable parameters*. Furthermore, this “model” based on chromatin states overcomes the defect of predicting negative recombination rates when gene density is high.

### The model with discrete chromatin states predicts fine-scale recombination patterns

Fig. 2B (bottom) shows the recombination rate pattern along genes and their 3 kb flanking regions (same syntenic genes and binning methodology as for the top of that figure). Regions just upstream of the TSS are richer in crossovers than regions downstream of the TTS which themselves are richer than gene bodies. Interestingly, these recombination patterns can be quite well *predicted* by the proportions of each chromatin state (top of Fig. 2B) using the experimentally measured state-specific recombination rates as displayed by the continuous blue curve in Fig. 2B (bottom).

We performed the analogous analysis on intergenic regions as shown in Fig. 2C (bottom). Again the experimental behavior is well predicted by our model that assigns one recombination rate to each chromatin state (cf. blue curve).

### Recombination rate is suppressed in small intergenic regions

The profiles and patterns in Fig. 2B and C pool gene bodies or intergenic regions, ignoring their sizes. To further test the model, we have considered the possibility that recombination rate patterns might vary as a function of the size of the region. For instance, the content in exons and introns is quite different for small and large genes and so this could potentially affect recombination rates.

To study the possible influence of gene body size, we divided the genes into size quantiles and recalculated the corresponding state occurrence profiles and recombination rate patterns. As illustrated in Supplementary Fig. S3, gene body size strongly affects chromatin state content. Furthermore, recombination rate patterns become more contrasted as gene size increases, with a concomitant decrease in the average recombination rate. Nevertheless, the model of 10 chromatin states correctly predicts these trends as shown by the blue curves.

The analogous study for intergenic region size is summarized in Supplementary Fig. S4, S5 and S6, treating separately the three possible orientations of the genes flanking the intergenic region: divergent, convergent and parallel. In contrast to the gene body case, the 10 chromatin state model’s predictions (blue curves) are not so good: the model significantly over-estimates the recombination rates when the size of the intergenic region is small.

To quantify this result, consider how the *average* recombination rate within intergenic regions depends on region size. In Fig. 3 we display this dependence, for all intergenic regions pooled (top) or separated according to the orientation of their flanking genes (bottom). There is a clear suppression of recombination rate when the size of the intergenic regions is less than 1.5 kb, while beyond 2.5 kb the curves are rather flat, with perhaps a trend to decrease beyond 10 kb. Fig. 3 also displays the recombination rates *predicted* when using the 10 state chromatin model. Clearly, the predictions over-estimate the recombination rate when the size of intergenic regions is small, in agreement with the trends seen in Supplementary Fig. S4, S5 and S6.

**Figure 3.**
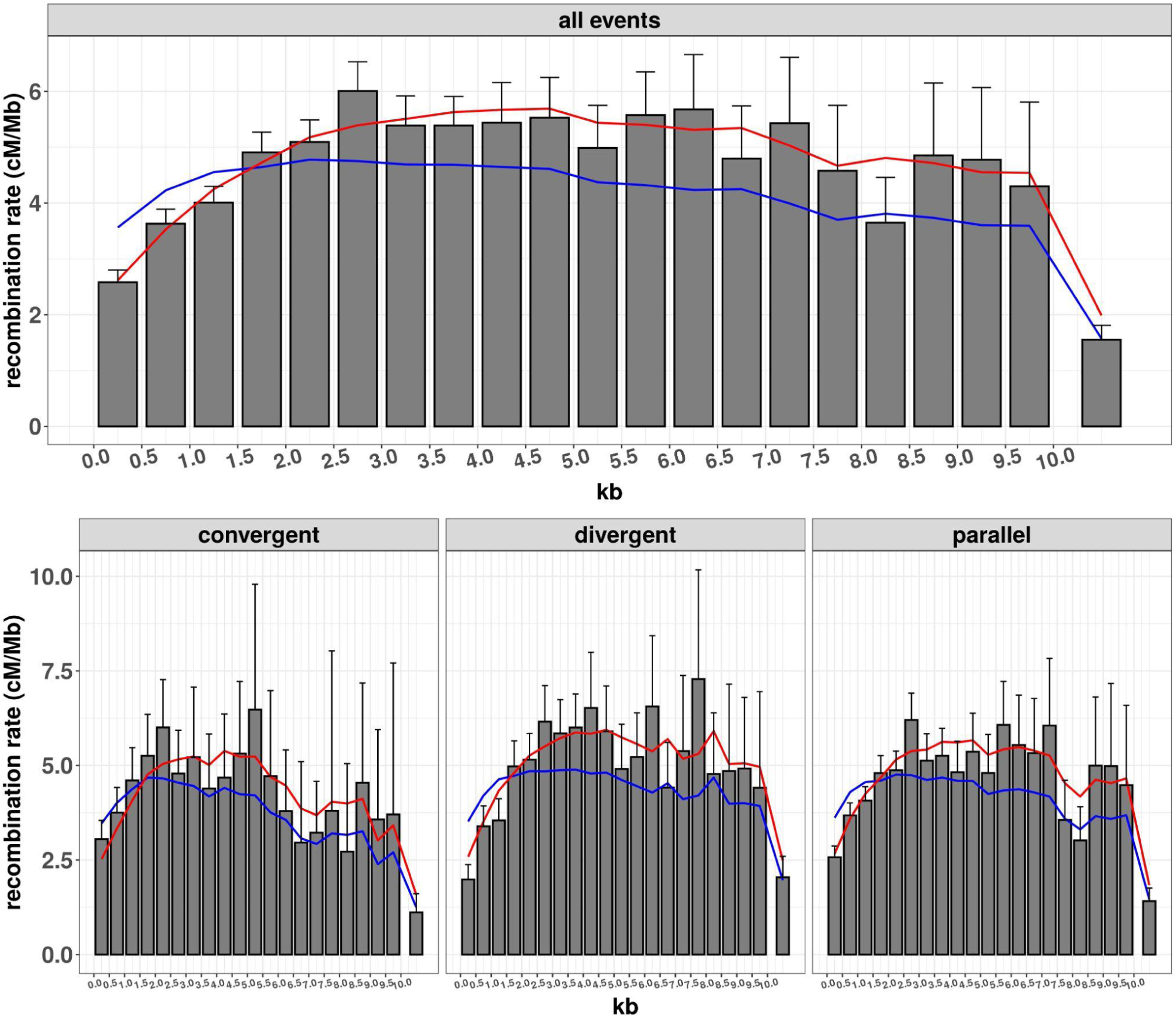
The relationship between the size of intergenic regions and their average recombination rate. These bar charts were constructed using all intergenic regions, but in the bottom the regions were divided into three categories according to the transcription orientations of the two flanking genes, corresponding to convergent, divergent and parallel transcriptions. In all cases, the X axis gives the size of the intergenic regions in kb, and the Y axis gives the corresponding averaged recombination rate (cM/Mb). Binning of the intergenic region sizes was applied every 500 bases up to a total size of 10 kb. For example, the leftmost bin covers intergenic regions of size 0 to 0.5 kb. However we also include a rightmost bar on each chart to cover intergenic regions of sizes larger than 10 kb. Error bars are errors on the mean computed by the jackknife method (only the top segments are displayed). In both top and bottom figures, the blue curves give the predicted recombination rates using the genome-wide recombination rates of the 10 chromatin states as obtained from Fig. 2A. The red curves show the predicted recombination rates when one includes the modulation based on the size of the intergenic regions as specified in Eq. 4.

These results motivated us to improve the model by including a modulation factor taking into account the sizes of intergenic regions. We parameterize such an effect by multiplying the recombination rate *r*_*i*_ of a segment in state *i* by the factor

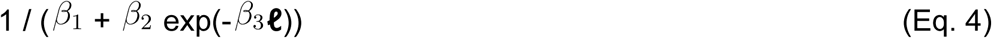

whenever the segment lies within an intergenic region of size ***ℓ*** kb. The detailed form of this modulation function is not so important but it should go smoothly from its minimum at ***ℓ*** =0 to its maximum at large ***ℓ***. The quantities *β*_1_, *β*_2_ and *β*_3_ are free parameters that we can adjust to minimize the deviation between observed and predicted recombination rates over all intergenic regions. The red curves in Fig. 3 show the corresponding improved predictions when including this modulation effect.

### Recombination rate is suppressed in regions of low SNP density

A high divergence between homologs suppresses recombination rate, a trend that is visible in the top left of Fig. 4 where SNP density is used as a proxy for divergence between homologs. However we see that *low* SNP density is also associated with reduced recombination. To confirm that this is not an artefact of the Rowan *et al*. (2019) dataset, we examined 5 other crosses published by Blackwell *et al*. (2020) who had found the same effect. The minor differences between our panels and those in their paper come from using different choices in the analysis pipelines: including or not the pericentromeric regions, using a bin size of 100 kb vs of 1 Mb, applying different filtering criteria to the remapped reads to define SNPs, and forbidding or not the fitting function to have negative values. The important point is that the two independent analyses reach the same conclusion: low SNP density is associated with lower recombination rate.

**Figure 4.**
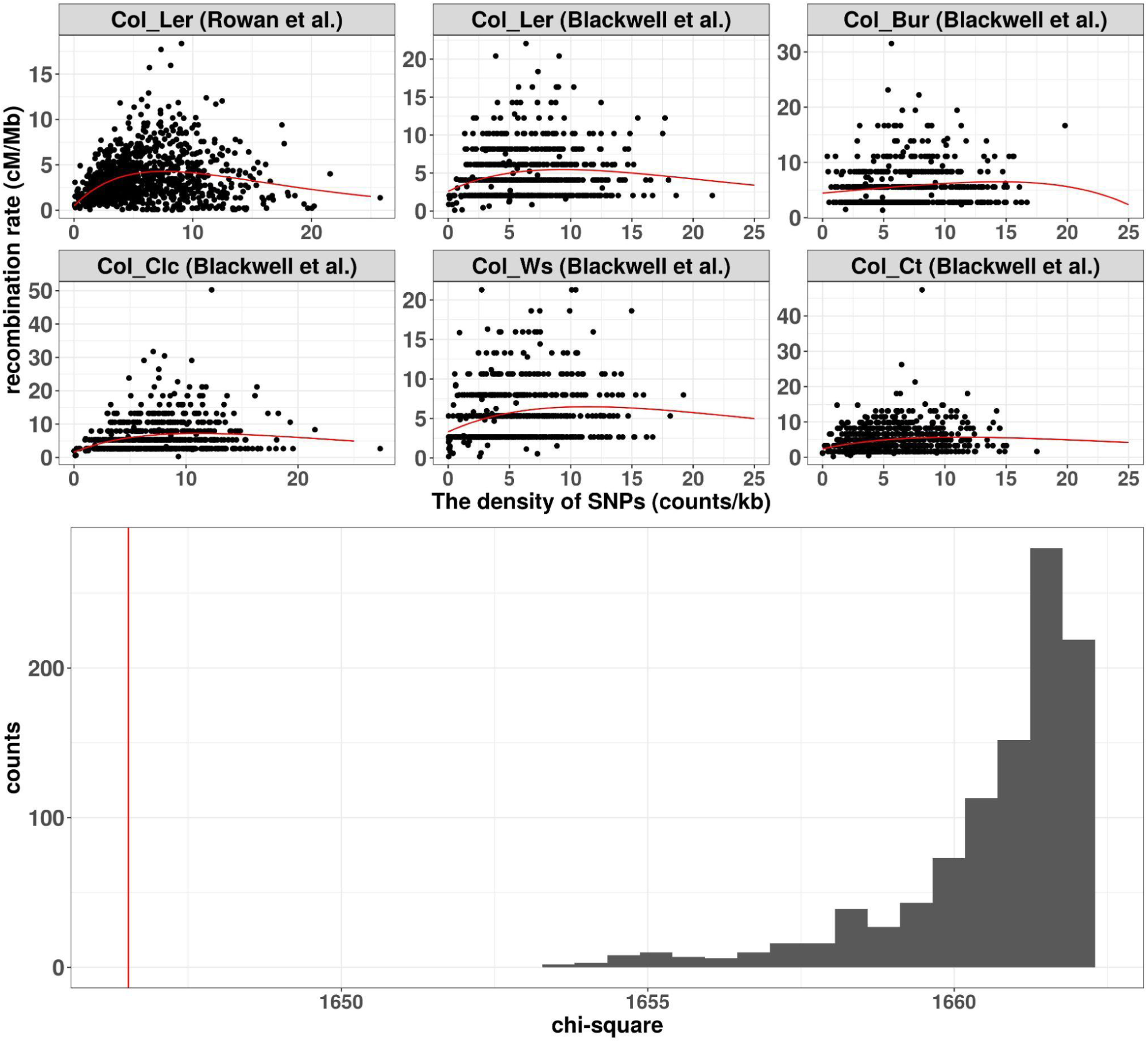
The relationship between recombination rate and SNP density. Top: The Col-0 genome was decomposed into bins of 100 kb. For each cross starting with that of Rowan et al. (2019), SNPs and COs were inferred from reads produced from the F2 populations by mapping to the Col-0 genome. SNP density and recombination rate were then determined for each bin and displayed as a scatter plot. The 5 additional crosses are from Blackwell et al. (2020). The continuous red curves are fits when using the function (a + b x) exp(-cx) based on minimizing the chi-square. All crosses show a reduced recombination rate at low SNP density. Bottom: Low SNP density is likely causally-responsible for reduced recombination rate. The hypothesis of no causal relation within the first two quantiles of SNP density is tested by shuffling the SNP content between the 5 F2 populations of Blackwell et al. (2020). X axis is chi-square (measuring goodness of fit) when allowing a position-dependent reference recombination rate to be modulated by SNP density using the same function as above. Y axis gives counts of simulations with shuffled SNPs, these counts being proportional to the probability density of chi-square under H0. The red vertical line is the actual chi-square value in the experimental dataset (unshuffled SNPs), showing that the recombination rate modulation, when using the SNPs between the parents of each separate cross, improves the fit far more than expected by chance (p-value ≼ 0.001).

### Low SNP density may be a causal factor of recombination rate suppression

In natural populations undergoing panmictic reproduction and subject to spontaneous mutations, drift generates linkage disequilibrium depending on recombination rate. Indeed, if a region of the genome has lower than average recombination rate, it will sustain larger haplotypic blocs and so its SNP density will be below average, producing the kind of correlation found in the top of Fig. 4. However *Arabidopsis thaliana* is a selfer, so linkage disequilibrium and thus the pattern of accumulation of mutations should not be affected by recombination. Specifically, if we consider the most recent common ancestor to Col-0 and Ler, it produced two separate lineages by successive generations of selfings, lineages in which mutations have accumulated *independently*. Under such dynamics, recombination cannot influence SNP density unless recombination itself generates mutations. This last possibility has long been downplayed because homologous recombination was considered to be nearly error-free (Guirouilh-Barbat *et al*., 2014), but it is now known that CO formation produces mutations in human (Arbeithuber *et al*. 2015, Halldorsson *et al*., 2019). In the absence of any such evidence in plants, we formalized as follows a test for the possibility that SNP density influences recombination.

Our statistical test based on the 5 F2 populations provided by Blackwell *et al*. (2020). In our framework (see M&M), we compare two hypotheses, H0 and H1. Under H0, we assume that there is an (unknown) “reference” recombination landscape, likely driven by genomic or epigenomic features, but common to all crosses. Under H1, this common landscape is modulated by the divergence between the homologs present, thus differently in each cross. We confront H0 to H1 by asking whether a good fit to the data necessitates the suppressive effect at low SNP density. We thus compare the chi-square goodness of fit using H1 to what would be expected if there were no causal suppressive effect (the H0 hypothesis). In the bottom of Fig. 4 we display the distribution of chi-square values under H0. Also shown is the chi-square value obtained using H1. The conclusion is that H1 is favored over H0 with a *p*-value smaller than 0.001.

### A state-based quantitative model with multiple factors modulating recombination rate has good predictive power

Our quantitative model builds on the framework of 10 discrete chromatin states by assigning to each an adjustable base recombination rate but also by applying three context-dependent modulating factors. The first factor is associated with intergenic size ***ℓ***: we parameterize the multiplicative modulation *via* the function 1/(*β*_1_ + *β*_2_ exp(-*β*_3_**ℓ**)) where **ℓ** is the size of the intergenic region in kb. The second factor is associated with SNP density **ρ**: we multiply the recombination rate by the factor (1 + *α*_1_**ρ**) exp(-*β*_2_ **ρ**). Lastly, at the whole chromosome level, it is known that CO numbers are tightly regulated with the result that genetic lengths hardly vary with genome size, independently of chromatin states. This regulation arises through both CO “interference” (COs tend to be well separated) and the obligatory CO (there is at least one CO per bivalent). As a result, the recombination rate of a specific genomic segment will be significantly higher if it belongs to a small chromosome than if it belongs to a large one. To incorporate this chromosome-wide effect, we rescale all predicted recombination rates within a chromosome to enforce its experimentally measured genetic length.

Overall our model has 15 adjustable parameters: the 10 base recombination rates and the 5 additional parameters for the modulation effects (the chromosome-specific rescalings do not require introducing any parameters or fits). To calibrate the resulting quantitative model, we compute a sum of squares which quantifies the deviation between the model’s predicted rates and the experimental ones from Rowan *et al*. (2019) when using a binning along the genome (see M&M for details). The optimized parameters are provided in Supplementary Table S3 when calibrating the model over the whole genome using various bin sizes. In Supplementary Fig. S7 we compare the predictions of recombination rate in our quantitative model to the experimental ones when using bins sizes ranging from 50 to 500 kb. One can also do the comparison at the level of the recombination landscapes: in Fig. 5 we show the predicted and experimental landscapes for chromosome 1 when using bins of size 100 kb (cf. Supplementary Fig. S8 for the other chromosomes). We see that the adjusted model reproduces much of the qualitative structure of the landscape. The inset in Fig. 5 provides a zoom on a region in the right arm, allowing one to better see the small scale trends. Even for this bin size which is rather large compared to the typical distance between genes, the model and experimental landscapes are far from smooth. Furthermore, both in the inset and in the main part of the figure we see that though there is quite a lot of concordance between the two curves for local minima and maxima, the model’s landscape generally underestimates the observed variance. This is partly due to the experimental landscape being subject to the stochasticity of CO numbers but it may also point to other determinants that could be missing in our analysis or data.

**Figure 5.**
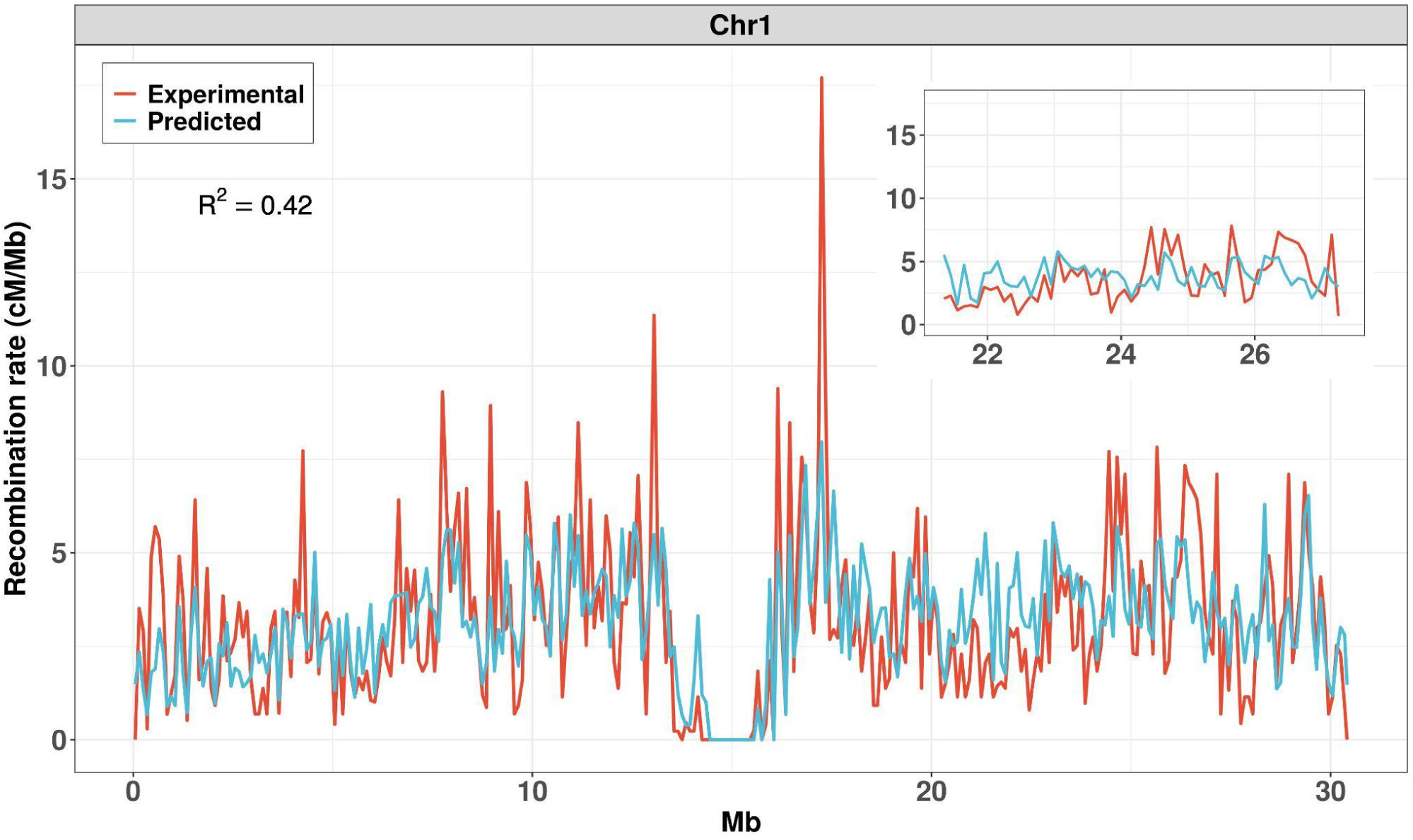
Experimental and predicted recombination landscapes of chromosome 1. Landscapes using 100 kb bins obtained from the Rowan *et al*. dataset (red) and predicted from our calibrated model based on chromatin states (blue). Inset: a zoom in the right arm. For landscapes of all chromosomes, see Supplementary Fig. S8.

Finally, to test the *predictive power* of our modeling approach and ensure that it does not introduce overfitting, we have calibrated the model on one chromosome and then used that calibration to predict recombination on the other chromosomes. Supplementary Table S4 gives the corresponding values of R^2^. For comparison, we perform the same test in Supplementary Tables S5 and S6 when using the additive model (Eq. 1) or its extension with interactions (Eq. 3). Clearly our model has significantly higher predictive power than those other models.

## Discussion and conclusions

### Aggregated chromatin states as predictors of recombination rate

The genome-wide distribution of COs is expected to follow largely from the degree to which the double strand break machinery can access the DNA. This will depend of course on the state of the chromatin and indeed many genomic and epigenomic features are empirically found to correlate with recombination rate. Unfortunately, the associated relations are typically non monotonic as displayed in Fig. 1. As a result, recombination rate modeling using these features as quantitative factors requires non linearities and leads to an unmanageable combinatorial complexity (cf. the 46 parameters in Eq. 3), not to mention problems for interpreting the resulting models and low prediction power(cf. Supplementary Tables S5 and S6). To overcome this difficulty, we use a classifier approach to automatically aggregate 16 genomic and epigenomic features into discrete classes (Sequeira-Mendes *et al*., 2014). This defines the starting point of our modeling wherein each position of the genome is considered to be in one of 10 chromatin states. Using the genome-wide recombination rates in each of these 10 states, Figs. 2B,C show that recombination patterns around genes and in intergenic regions are rather well predicted at a qualitative level. In particular, near the ends of genes, this simple modeling leads to enhanced recombination rates, in agreement with experiment (Choi *et al*., 2013; Kianian *et al*., 2018; Marand *et al*., 2017).

### Intergenic region size modulates recombination rate

The simple model using genome-wide recombination rates in each of the 10 states does not adequately predict the suppressed recombination rate in small intergenic regions (cf. Fig. 3). This suppression effect could be the consequence of a local context affecting chromatin accessibility for biophysical reasons. A first such reason could be that small intergenic regions are partly hidden from the double strand break machinery by their flanking regions when these are in dense chromatin. A second such reason could be the way chromatin loops are organised in meiosis; if denser chromatin (e.g. containing gene bodies) is preferentially tethered to the base of those loops, it will pull along with it adjacent stretches of open chromatin, hiding these from the double strand break machinery.

### Lack of any sequence divergence may drive lower recombination rate

The empirical data in multiple crosses show that regions with very low divergence between homologs typically have low recombination rate (cf. Fig. 4). That is expected in panmictic populations where recombination shapes linkage disequilibrium and thus SNP density. However *A. thaliana* is a selfing species with a very low rate of outcrossing of about 2% (Hoffmann *et al*., 2003; Platt *et al*., 2010). That leads to low genetic divergence within given habitats which is further exacerbated by adaptive pressures, so recombination in the wild will hardly do any allelic shuffling. We thus argue that the data in this species might be explained if an absence of divergence between homologs causally suppresses COs. Clearly, such an effect makes sense from an evolutionary perspective: if a genomic region has no underlying sequence diversity, there is little point in producing COs there.

Interestingly, a reduction of recombination rate caused by near perfect sequence homology was demonstrated by two previous works on *A. thaliana*. First, Ziolkowski *et al*. (2015) considered a heterozygous block within an otherwise homozygous chromosome and found that CO frequency was enhanced in the heterozygous region. Second, Blackwell *et al*. (2020) showed that *msh2*, a mutant of mismatch repair, redistributed COs towards regions of low SNP density, suggesting that, in wild type, perfect sequence homology represses CO formation. The behaviors found in both of these works can be interpreted as the large-scale manifestation of the causal SNP effect we hypothesize.

### A quantitative model of recombination rate with good predictive power

Our full model integrates local genomic and epigenomic features but also context-dependent information. All of its 15 parameters have very direct interpretations. It has good predictive power as shown in Supplementary Tables S4 to S6 and is able to reproduce much of the variation in rates arising in the recombination landscape (cf. Fig. 5 and Supplementary Fig. S8). Clearly not all of the variation is captured by our model. First, there is statistical noise inherent to the experimental landscape. Second, although the model predicts major peaks and troughs in the landscape, it tends to underestimate their amplitude. This may suggest a form of competition between sites for recruiting the machinery that produces double strand breaks. There are also other caveats to our modeling. The most obvious one is that because of lack of appropriate data, we had to use measurements of epigenetic marks in Col-0 only and from tissues such as leaf or root rather than from meiocytes. Fortunately, it seems that the epigenetic landscape is largely shared between somatic and germline tissues, the differences being restricted to a small fraction of the genome (Walker *et al*., 2018). We did a systematic investigation of this point using published data (cf. Fig. S1) and showed that the epigenomic patterns are surprisingly similar between somatic and germline tissues. Another limitation of our modeling is that it necessarily ignores any sex-dependent differences in recombination landscapes, focusing only on the female-male average. Similarly, we have not included CO interference, another factor that shapes recombination landscapes. Lastly but perhaps very importantly, we take no account of the well known fact that meiotic chromosomes are organised in loops tethered to an axis. This structural aspect of meiotic chromosomes may be important for modulating recombination rates and it is tempting to conjecture that these loops may be responsible for the large peaks seen in the recombination landscape (cf. Fig. 5 and Supplementary Fig. S8). Unfortunately, very little is known about these loops, in particular concerning their size, position and variability across genetic backgrounds. Hopefully these uncertainties will be lifted in the near future, given that standard chromosome conformation capture techniques applied to meiotic cells should provide the required information quite directly.

## Acknowledgements

We thank T. Blein, M. Rousseau-Gueutin and P. Sourdille for discussions and we are particularly indebted to M. Grelon who provided feedback on our drafts. We are also grateful to B. Rowan and Ian Henderson for sharing their data.

## Author Contributions

MF and OM conceived the study. OM designed the modeling. YMH performed all the bioinformatics and code writing necessary for the subsequent analyses. All authors worked on the analyses and wrote the article.

## Conflicts of Interest declarations in manuscripts

None

## Data and Coding Availability Statement

This work produced no new data but exploited previously published data (see Material and Methods section, Supplementary Material and Supplementary Table S1). All pipelines and analysis codes are given as Supplementary files.

