## Supplementary_materials for "Quantitative modeling of fine-scale variations in the *Arabidopsis thaliana* crossover landscape"

| epigenomic features | Sample accession or series accession number | tissue | reference |
| --- | --- | --- | --- |
| H3K4me1 | GSM3674621 | leaves | Lu et al., 2019; Crisp et al., 2020 |
| | GSM4668649 | seedlings | Niu et al., 2021 |
| | GSM4609829 | root non- hair cells | missing |
| | GSM4785549 | inflorescence | Liu et al., 2021 |
| | E-MTAB-7370 | unopened flower buds | Lambing et al., 2020 |
| H3K4me3 | GSM3674620 | leaves | Lu et al., 2019; Crisp et al., 2020 |
| | GSM4154769 | seedlings | Liu et al., 2020 |
| | GSM2210857 | roots | Yen et al., 2017 |
| | GSM4785552 | inflorescence | Liu et al., 2021 |
|  | GSE120664 | sperm nuclei | Borg et al., 2020 |
| H3K9me2 | GSM4734580 | leaves | Wang et al., 2021 |
| | GSM3040062 | 10-day seedlings | Ma et al., 2018 |
| | GSM4422529 | mature embryos | Parent et al., 2021 |
| | GSM4818168 | flowers | Feng et al., 2020 |
| | E-MTAB-7370 | unopened flower buds | Lambing et al., 2020 |
| H3K27me3 | GSM3674617 | leaves | Lu et al., 2019; Crisp et al., 2020 |
| | GSM3617717 | seedlings | Shu et al., 2021 |

| | | | |
|---|---|---|---|
| | GSM2210865 | roots | Yen et al., 2017 |
| | GSM4785573 | inflorescences | Liu et al., 2021 |
| | GSE120664 | sperm nuclei | Borg et al., 2020 |
| ATAC | GSM3674715 | leaves | Lu et al., 2019; Crisp et al., 2020 |
| | GSM2719200 | stem cells | Sijacic et al., 2018 |
| | GSM2719204 | mesophyll cells | |
| | GSM3498708 | flowers | Potok et al., 2019 |
| | GSE155344 | microspores | Borg et al., 2021 |
| DNase | GSM1289358 | seedlings | Sullivan et al., 2014; Sullivan et al., 2019 |
| | GSM1289374 | whole roots | |
| | GSM1289378 | seed coats | |
| | GSM1289380 | open flowers | |
| | GSM1289381 | unopened flower | |

Supplementary Table S1. Origin and description of datasets for the 6 epigenomic features used in this study.

| | intercept (a_0) | gene (a_1) | TE (a_2) | TSS (a_3) | H3K4me1 (a_4) | H3K4me3 (a_5) | H3K9me2 (a_6) | H3K27me3 (a_7) | ATAC (a_8) | DNase (a_9) | $R^2$ |
|---|---|---|---|---|---|---|---|---|---|---|---|
| 50kb | 1.56** | -3.6*** | -1.67** | 0.17 | -0.04 | 0.05 | -0.004*** | 0.11*** | 0.65*** | 0.006* | 0.28 |
| 100kb | 1.00 | -5.02*** | -1.14 | 0.26 | -0.07 | 0.16* | -0.01*** | 0.14** | 0.71*** | -0.005 | 0.36 |
| 200kb | 0.06 | -4.44* | 0.23 | 0.3 | -0.09 | 0.16 | -0.01** | 0.14 | 0.75*** | -0.0007 | 0.42 |
| 500kb | -1.01 | -5.82 | 1.08 | 0.27 | -0.08 | 0.24 | -0.01 | 0.16 | 0.83*** | 0.003 | 0.50 |

Supplementary Table S2. Adjusted parameters and $R^2$ values for the additive model when using different bin sizes. The 9 successive features are those in Fig. 1 (ordered left to right and top to bottom). Parameter values were obtained using the lm() function in R. *, ** and *** correspond to parameters having *p*-values less than 0.05, 0.01 and 0.001 respectively for the hypothesis that the true value of the parameter vanishes. The first column gives the bin size used for each fit. Note that the statistical noise intrinsic to CO formation inevitably drives $R^2$ (last column) upward as bin size increases.

| name | 50k | 100k | 200k | 500k |
|---|---|---|---|---|
| r_state1 | 1.340778 | 1.621979 | 1.178138 | 0.576624 |
| r_state2 | 1.329113 | 1.648426 | 2.25995 | 1.526519 |
| r_state3 | 4.16E-09 | 2.75E-11 | 7.35E-12 | 1.42E-12 |
| r_state4 | 1.4614 | 1.673411 | 2.002888 | 1.189302 |
| r_state5 | 0.523589 | 0.64217 | 1.133385 | 0.418807 |
| r_state6 | 0.396265 | 0.05914 | 7.35E-12 | 1.42E-12 |
| r_state7 | 4.16E-09 | 2.75E-11 | 7.35E-12 | 1.42E-12 |
| r_state8 | 1.147018 | 1.539397 | 2.328571 | 1.228594 |
| r_state9 | 4.16E-09 | 2.75E-11 | 7.35E-12 | 1.42E-12 |
| r_SV | 4.16E-09 | 2.75E-11 | 7.35E-12 | 1.42E-12 |
| $\alpha_1$ | 1.186288 | 1.021344 | 1.027258 | 1.567372 |
| $\alpha_2$ | 0.086961 | 0.084894 | 0.0947 | 0.090547 |
| $\beta_1$ | 0.43086 | 0.429626 | 0.525843 | 0.395678 |
| $\beta_2$ | 83.79787 | 88207.18 | 127686.6 | 299934.3 |
| $\beta_3$ | 6.090687 | 9.646168 | 8.910464 | 9.705455 |
| $R^2$ | 0.40771 | 0.493876 | 0.569399 | 0.667817 |

Supplementary Table S3. Results of calibrating the quantitative model having 15 parameters when using bin sizes from 50 k to 500 k. In the column "name", r_state1 to r_SV refer to the "base recombination rate" for each of the 10 chromatin states, $\alpha_1$ and $\alpha_2$ (respectively $\beta_1$, $\beta_2$. $\beta_3$) refer to the parameters in the SNP (respectively intergenic-region size) modulation factor, and finally $R^2$ refers to the fraction of the variance explained by the model (cf. Eq. 2).

| | Chr1 (fit) | Chr2 (fit) | Chr3 (fit) | Chr4 (fit) | Chr5 (fit) |
|---|---|---|---|---|---|
| Chr1 (predict) | 0.469 | 0.293 | 0.341 | 0.243 | 0.444 |
| Chr2 (predict) | 0.406 | 0.514 | 0.469 | 0.440 | 0.447 |
| Chr3 (predict) | 0.526 | 0.553 | 0.615 | 0.461 | 0.560 |
| Chr4 (predict) | 0.433 | 0.471 | 0.482 | 0.552 | 0.462 |
| Chr5 (predict) | 0.450 | 0.348 | 0.395 | 0.318 | 0.477 |

Supplementary Table S4. Goodness of fit and predictive power of the model with 15 parameters. Displayed are the $R^2$ values when using one chromosome (labeling columns) to fit the model and applying that calibration to predict recombination rates on all chromosomes. The genome has been segmented into bins of size 100 kb. Note that in each row the largest $R^2$ value must occur for the chromosome that has been used to do the adjustment of parameters. Omitting the $R^2$ values produced by the calibrations (on the diagonal), the average $R^2$ of the predictions (remaining 20 values) is 0.427.

| | Chr1 (fit) | Chr2 (fit) | Chr3 (fit) | Chr4 (fit) | Chr5 (fit) |
|---|---|---|---|---|---|
| Chr1 (predict) | 0.348 | 0.222 | 0.22 | 0.171 | 0.292 |
| Chr2 (predict) | 0.211 | 0.409 | 0.263 | 0.344 | 0.339 |
| Chr3 (predict) | 0.138 | 0.35 | 0.455 | 0.353 | 0.383 |
| Chr4 (predict) | 0.218 | 0.347 | 0.34 | 0.383 | 0.328 |
| Chr5 (predict) | 0.281 | 0.218 | 0.274 | 0.215 | 0.346 |

Supplementary Table S5. Goodness of fit and predictive power of the additive model (Eq. 1) with 10 parameters exploiting the genomic and epigenomic features of Fig. 1. Displayed are the $R^2$ values when using one chromosome (labeling columns) to fit the model and applying that calibration to predict recombination rates on all chromosomes (same procedure as in Supplementary Table S4, again with bins of size 100 kb). Omitting the $R^2$ values produced by the calibrations (on the diagonal), the average $R^2$ of the predictions (remaining 20 values) is 0.275.

| | Chr1_fit | Chr2_fit | Chr3_fit | Chr4_fit | Chr5_fit |
|---|---|---|---|---|---|
| Chr1_predict | 0.447 | -1.364 | -0.493 | -0.407 | 0.176 |
| Chr2_predict | -0.299 | 0.579 | -0.614 | -39.286 | -8.073 |
| Chr3_predict | -0.307 | -78.667 | 0.568 | -39.829 | -2.22 |
| Chr4_predict | 0.074 | -17.86 | 0.001 | 0.545 | -0.349 |
| Chr5_predict | -0.3 | -27.968 | -1.393 | -2.783 | 0.501 |

Supplementary Table S6. Goodness of fit and predictive power of the model with interactions (Eq. 3) with 46 parameters exploiting the genomic and epigenomic features of Fig. 1. Displayed are the $R^2$ values when using one chromosome (labeling columns) to fit the model and applying that calibration to predict recombination rates on all chromosomes (same procedure as in Supplementary Table S4, again with bins of size 100 kb). Note that the $R^2$ of most of the predictions are negative, showing that this model with interactions has no predictive power, presumably because it strongly overfits the data during calibration.

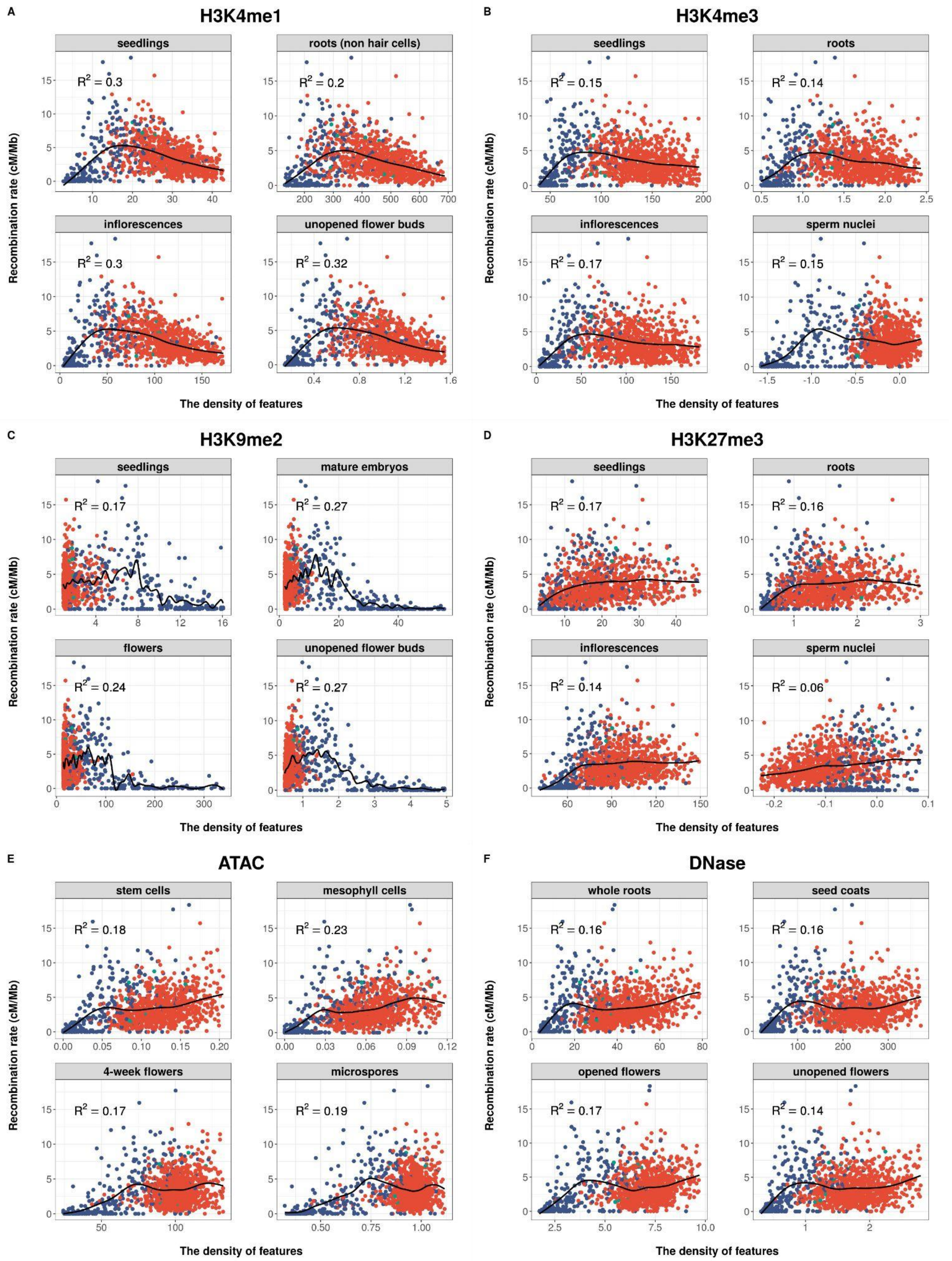


Supplementary Figure S1. The correlations between recombination rate and six epigenomic features when measured in somatic and germinal tissues. From (A) to (F), each sub figure combines four plots using data from two somatic and two germinal tissues for the same epigenomic feature. The subtitle on each plot indicates the data's accession number and the tissue name. Each dot represents the values for a 100-kb bin. The x-axis shows the density of peaks or reads of each feature according to the format of raw data downloaded from NCBI or ArrayExpress databases, and the y-axis is the recombination rate based on a total of 17,077 crossovers from the Col-0-Ler $F_2$ population. For display purposes, we truncated the 2.5 % extremities on both extremities of the x-axes.

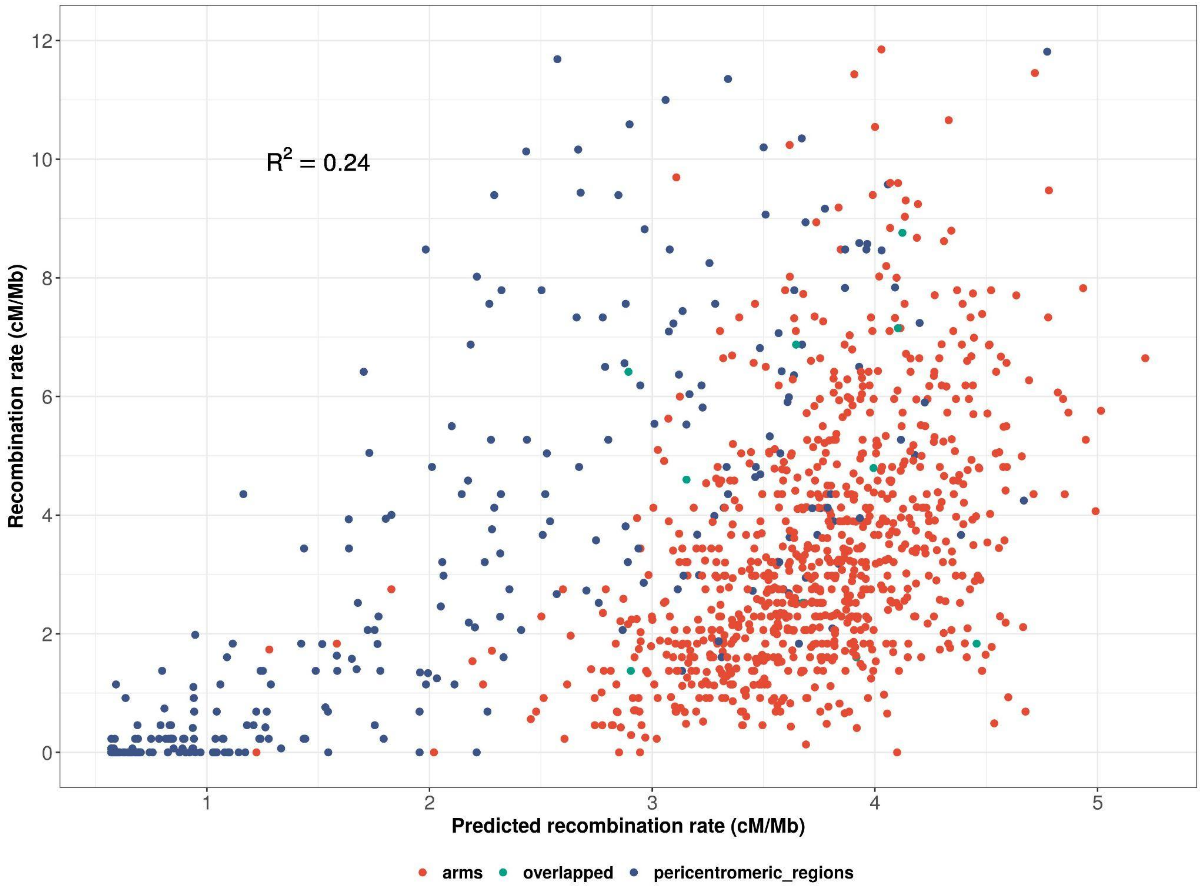


Supplementary Figure S2. Comparison of experimental and predicted recombination rates. Here the predictions are those of the 10 chromatin states model using the experimentally measured state-specific recombination rates (no adustable parameters). Each data point is associated with a bin of 100 kb along the genome. The fraction of variance explained by the model is $R^2$ = 0.24.

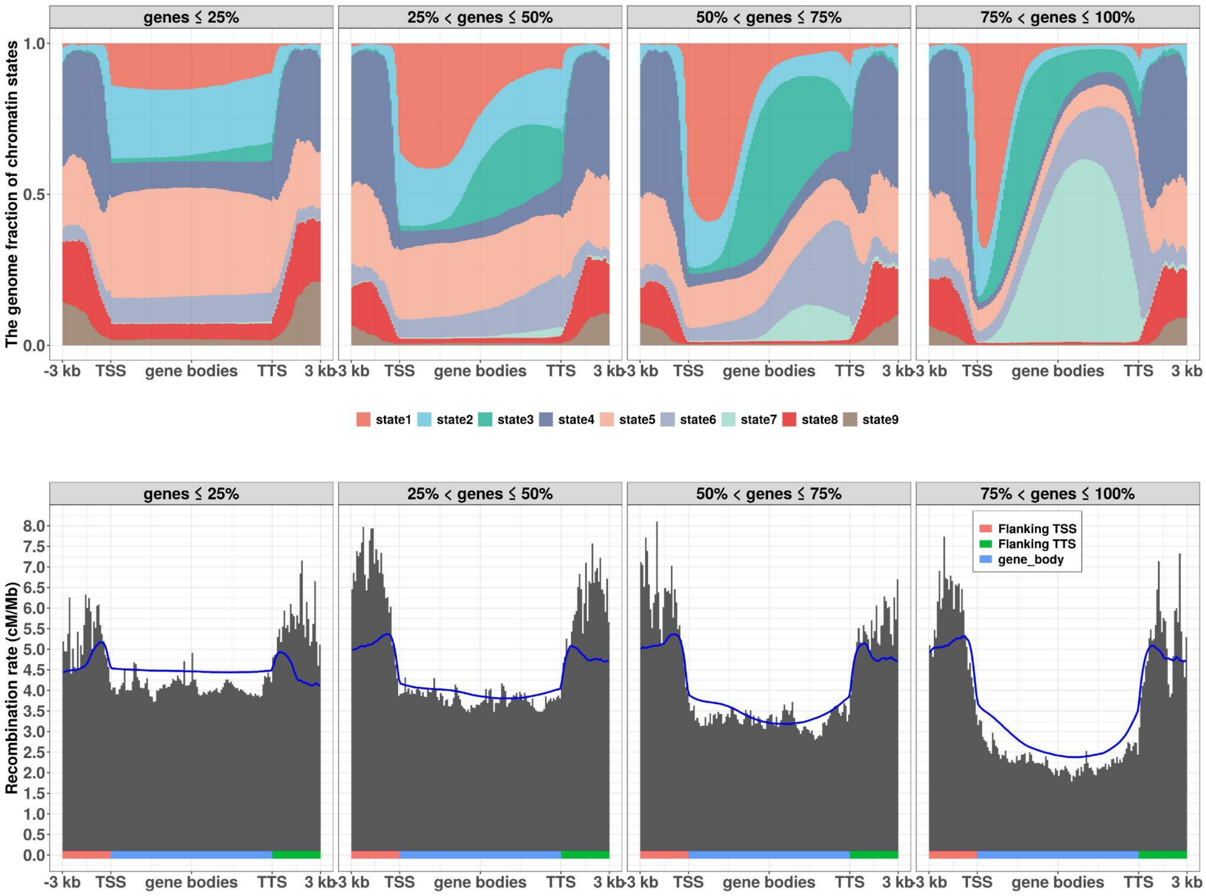


Supplementary Figure S3. Dependence of recombination patterns on gene body size. The profiles of chromatin states and the recombination rate patterns are determined separately in the four quantiles of gene size. The procedures are the same as in Fig 2B, and the blue curve shows the prediction of the model with 10 chromatin states when using the experimentally measured state-specific recombination rates (no adjustable parameters).

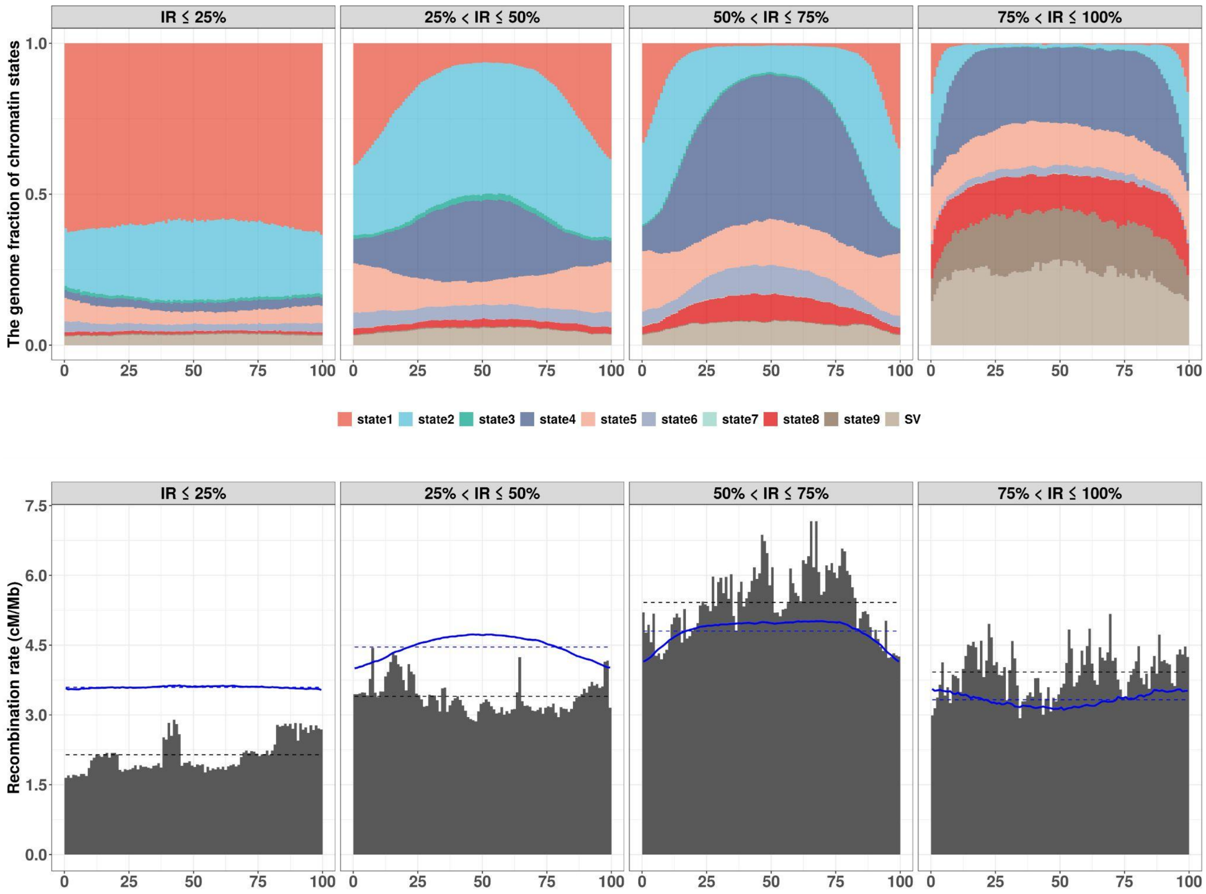


Supplementary Figure S4. The profiles of chromatin states and recombination rate in intergenic regions where their flanking genes are divergent orientation. All "divergent" intergenic regions larger than 100 base pairs are divided into 4 groups depending on their size, and each group has equal 25 % of intergenic-region events. In each group, we forced every intergenic region to be segmented into 100 bins, then pooled all data of each bin, and calculated the fraction of 9 chromatin states and SV and recombination rate of each bin. At the top of this figure, it shows the fraction of states on the y-axis, and 100 bins on the x-axis. At the bottom, the y-axis is the recombination rate, and x-axis indicates bins. The bottom histograms show the experimental recombination rate of 100 bins, and the black dashed lines are the average CO rate of the experimental ones. The procedures are the same as in Fig 2B, and the blue curve shows the prediction of the model with 10 chromatin states when using the experimentally measured state-specific recombination rates. The blue dashed lines are the average predicted recombination rate (no adjustable parameters).

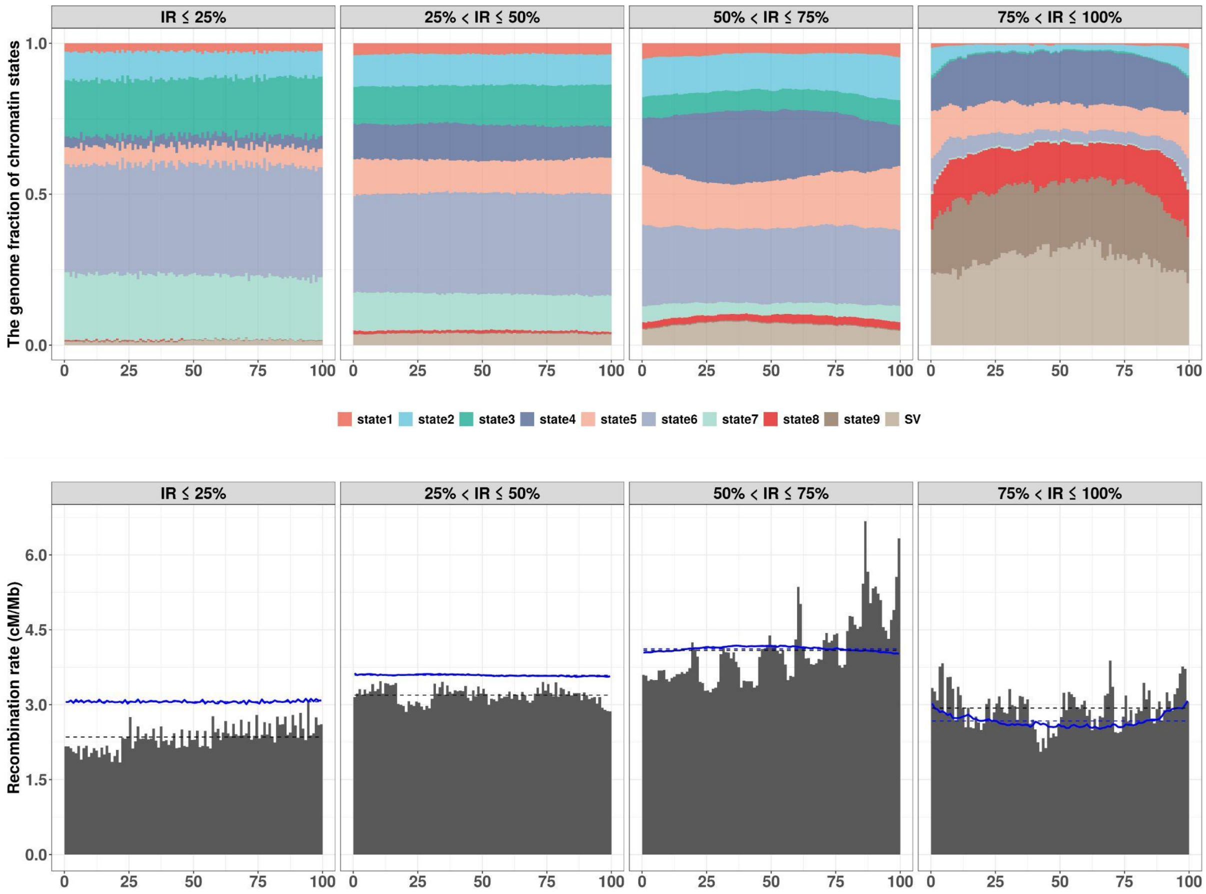


Supplementary Figure S5. The profiles of chromatin states and patterns of recombination rate in intergenic regions where their flanking genes have convergent orientation. The procedures and quantities displayed are the same as in Supplementary Figure S4.

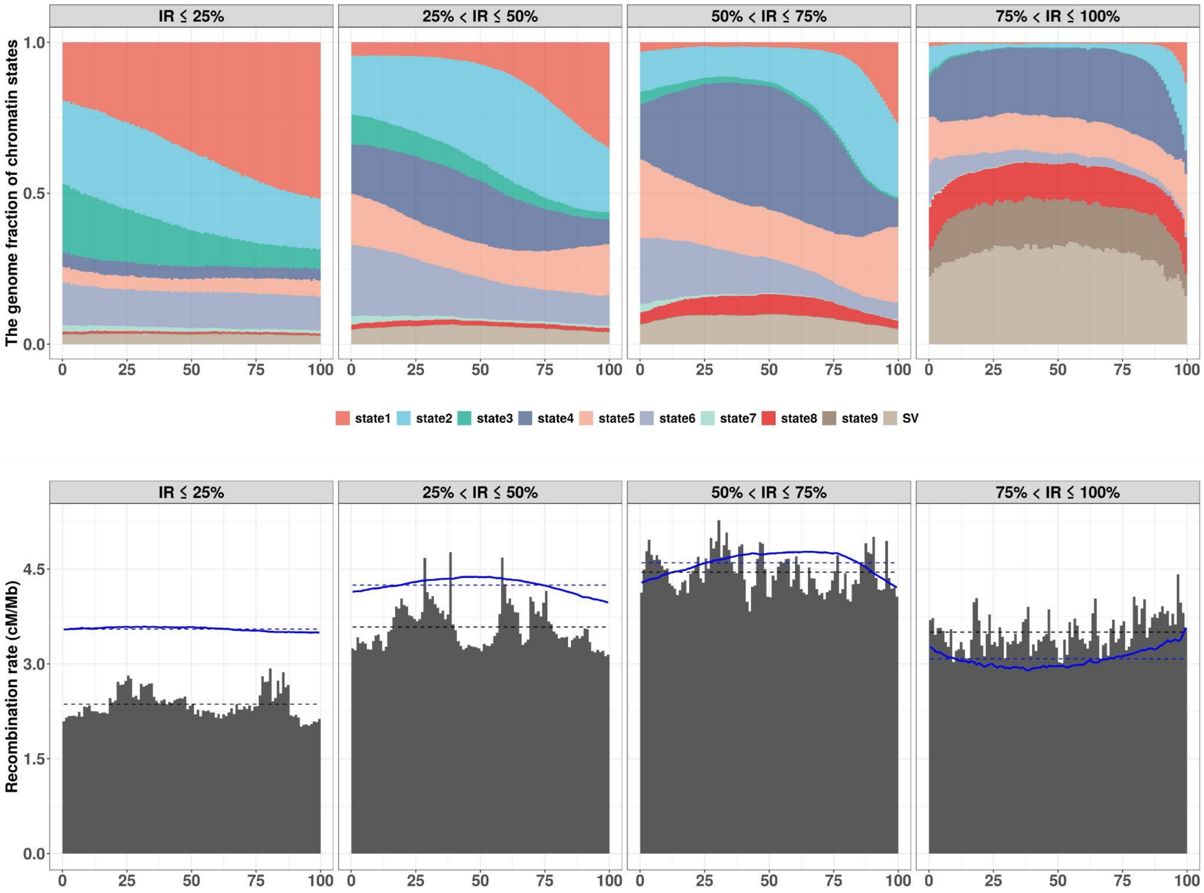


Supplementary Figure S6. The profiles of chromatin states and recombination rate in intergenic regions where their flanking genes are parallel orientation. The procedures and quantities displayed are the same as in Supplementary Figure S4.

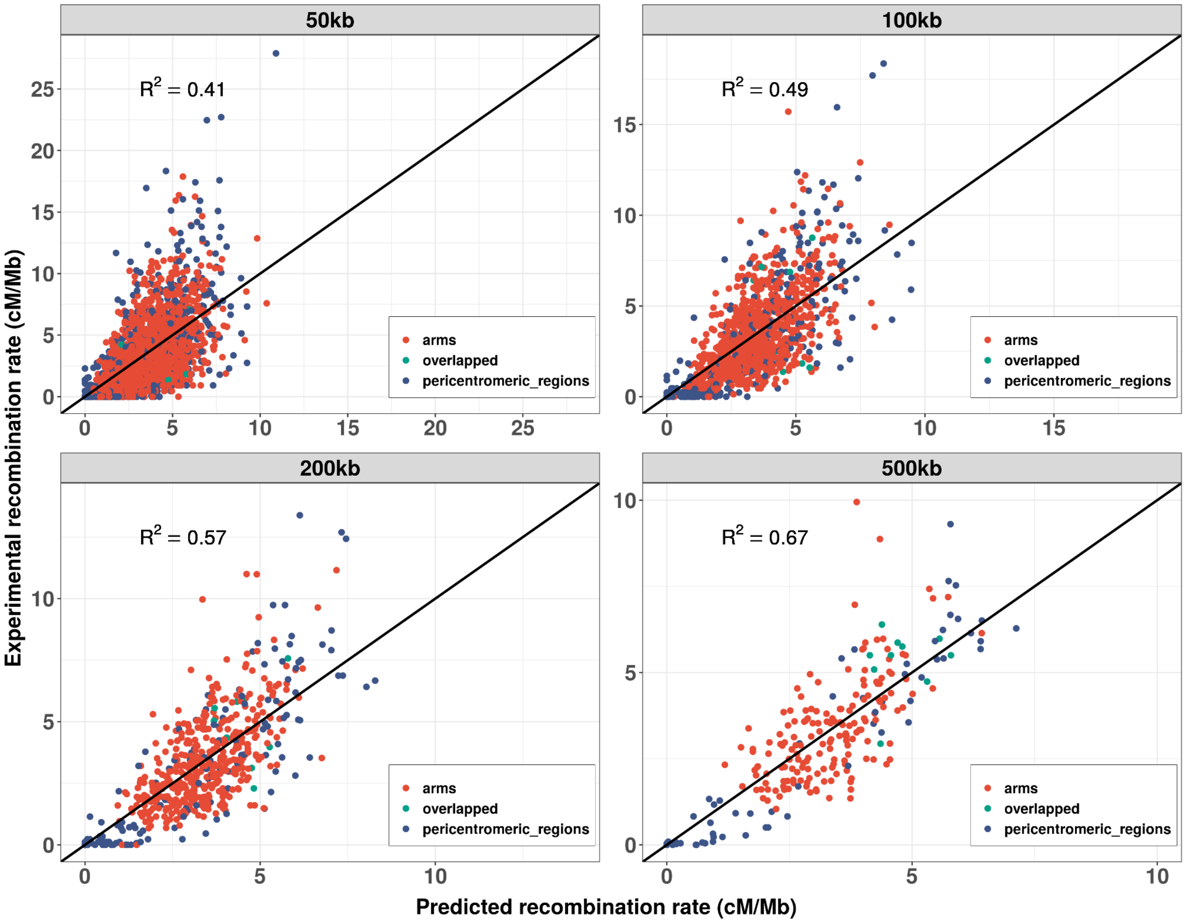


Supplementary Figure S7. Scatterplots of experimental and predicted recombination rate when the model calibration is done using bin sizes ranging from 50 to 500 kb. The X-axis is the recombination rate predicted by our quantitative model that incorporates 10 chromatin states along with contextual modulating factors, having a total of 15 adjustable parameters. The Y-axis is for the experimental recombination rate as produced from the Rowan *et al*. (2019) dataset. $R^2$ is the fraction of the variance explained by the model; it inevitably increases as bin size decreases because the CO numbers per Mb are more subject to stochastic noise.

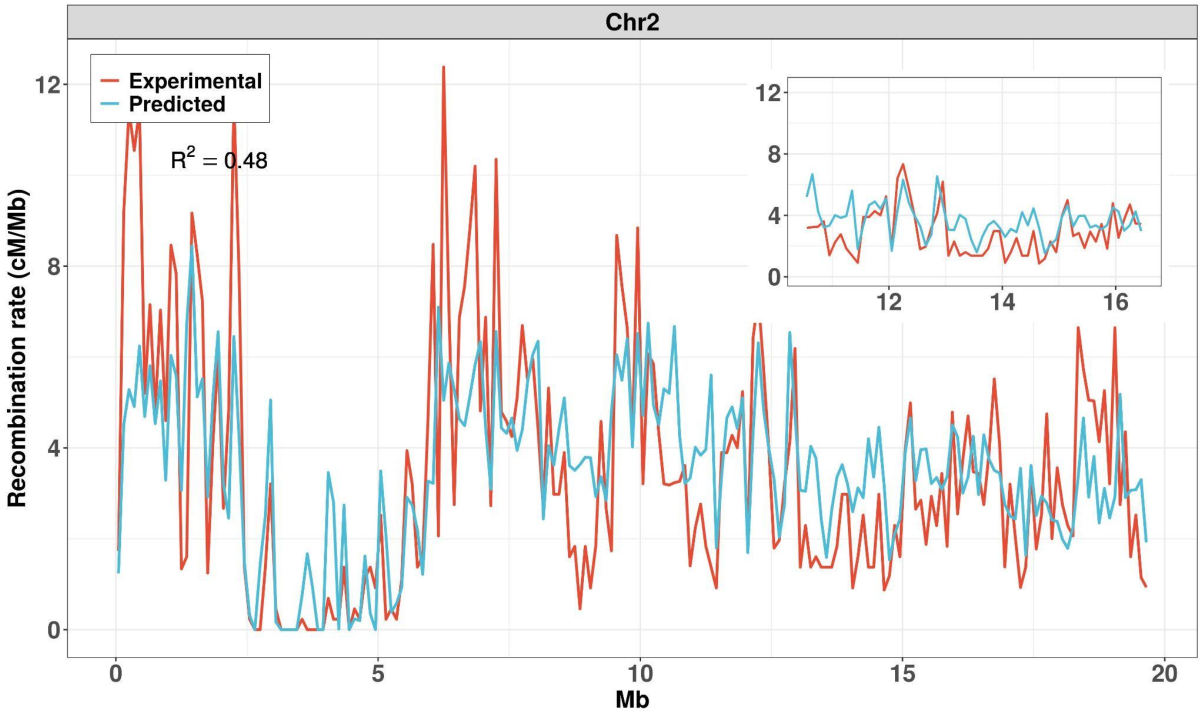
Chr2
Experimental
Predicted
$R^2 = 0.48$
Recombination rate (cM/Mb)
Mb

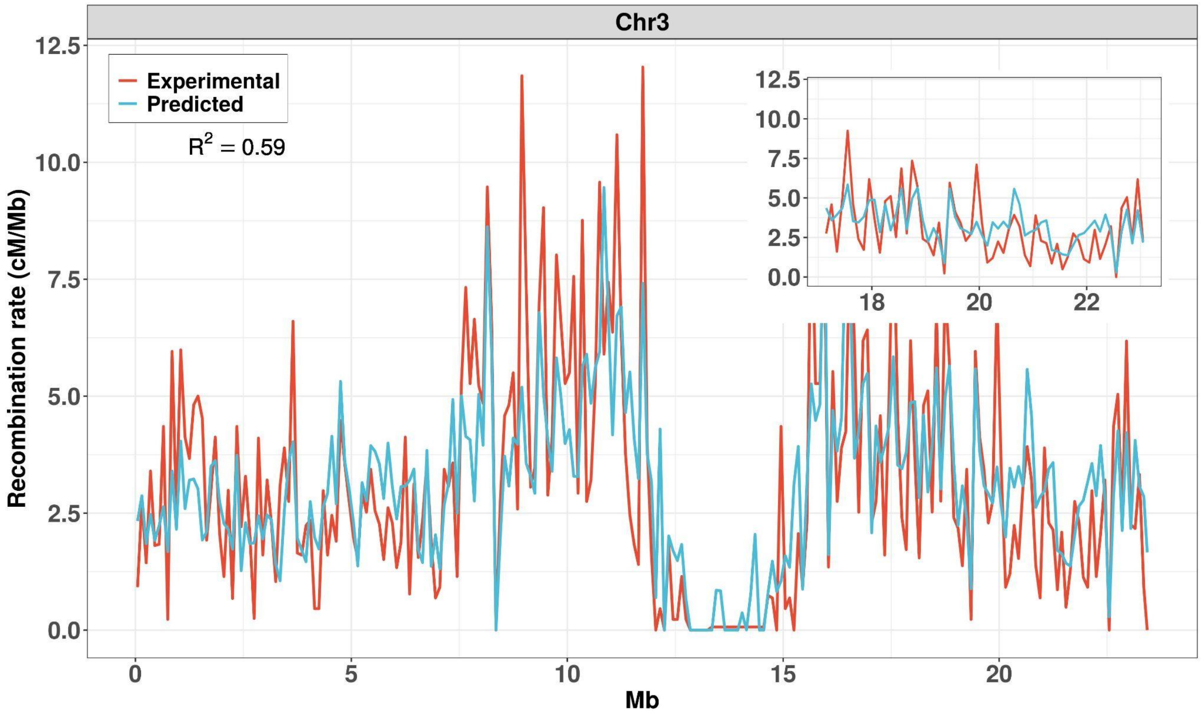
Chr3
Experimental
Predicted
$R^2 = 0.59$
Recombination rate (cM/Mb)
Mb

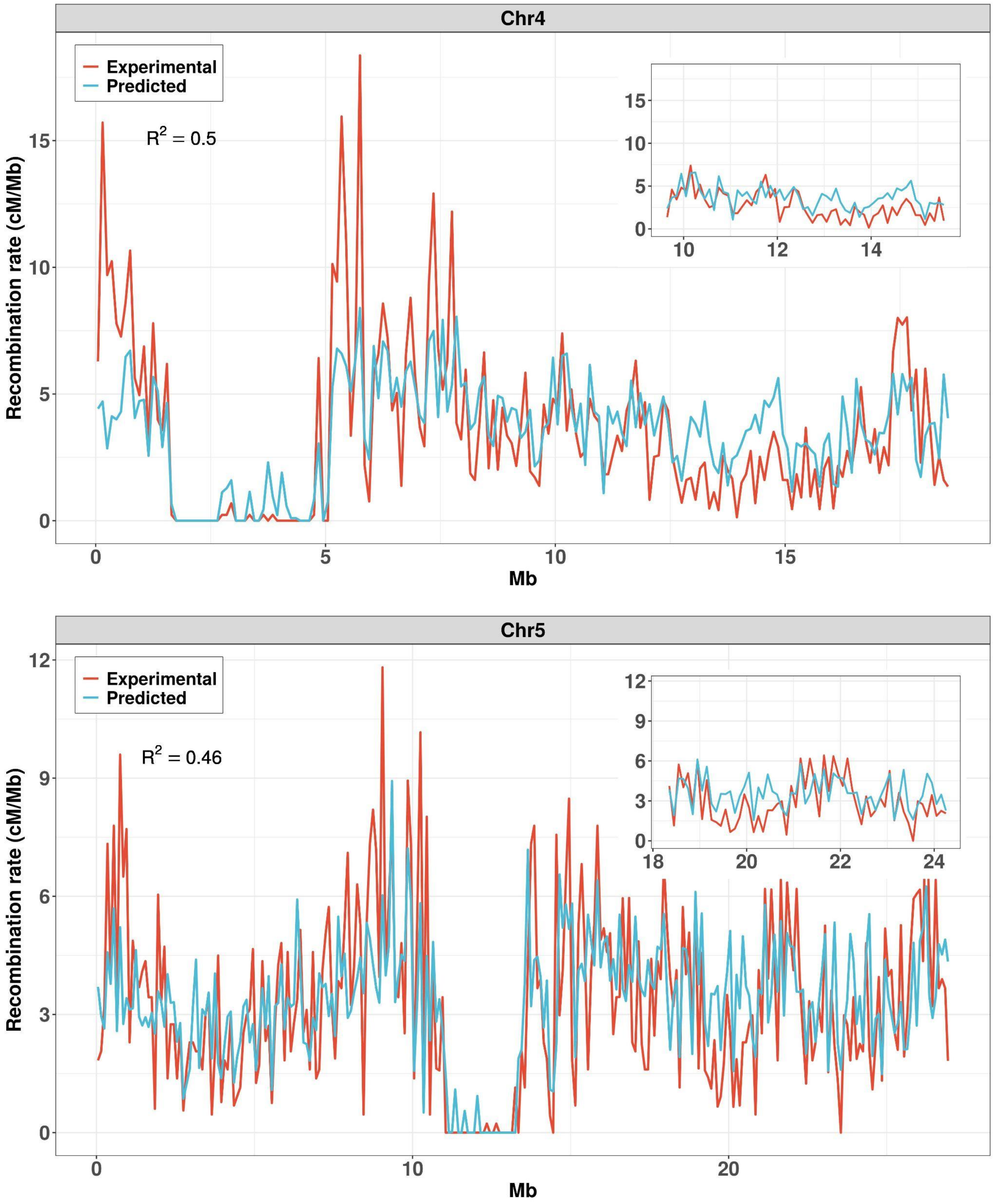


Supplementary Figure S8. Experimental and predicted recombination landscapes of chromosomes 2 to 5. Landscapes using 100 kb bins produced from the Rowan *et al*. dataset (red) and from our quantitative model with 15 adjustable parameters (blue). Inset: a corresponding zoom within the right arm. $R^2$ is the fraction of the recombination rate variance that is explained by the model.
